# Deep Learning and GBLUP Integration: An Approach that Identifies Nonlinear Genetic Relationships Between Traits

**DOI:** 10.1101/2024.03.23.585208

**Authors:** Fatima Shokor, Pascal Croiseau, Hugo Gangloff, Romain Saintilan, Thierry Tribout, Tristan Mary-Huard, Beatriz C.D. Cuyabano

## Abstract

**Background:** Genomic prediction aims to predict the breeding values of multiple complex traits, usually assumed to be normally distributed by the largely used statistical methods, thus imposing linear genetic correlations between traits. While statistical methods are of great value for genomic prediction, these methods do not account for nonlinear genetic relationships between traits. If such relationships exist, although statistical models do perform a fair linear approximation, their prediction accuracy is limited due to the nonlinearity. Deep learning (DL) is a promising methodology for predicting multiple complex traits, in scenarios where nonlinear genetic relationships are present, due to its capacity to capture complex and nonlinear patterns in large data. We proposed a novel hybrid DLGBLUP model which uses the output of the traditional GBLUP, and enhances its PGV by accounting for nonlinear genetic relationships between traits using DL. Using simulated data, we compared the accuracy of the PGV obtained with the proposed hybrid DLGBLUP model, a DL model, and the traditional GBLUP model – the latter being our baseline reference.

**Results:** We found that both DL and DLGBLUP models either outperformed GBLUP, or presented equally accurate PGV, with a particular greater accuracy for traits presenting a strongly characterized nonlinear genetic relationship. Overall, DLGBLUP presented the highest prediction accuracy, up to 0.2 points higher than GBLUP, and smallest mean squared error of the PGV for all traits. Additionally, we evolved a base population over seven generations and compared the genetic progress when selecting individuals based on the additive PGV obtained by either DL, DLGBLUP or GBLUP. For all traits with a nonlinear genetic relationship, after the fourth generation, the observed genetic gain when selection was based on the additive PGV from GBLUP was always inferior to the one achieved from either DL or DLGBLUP.

**Conclusions:** The integration of DL into genomic prediction enables the possibility of modeling nonlinear relationships between traits. Moreover, by identifying these nonlinear genetic relationships, our DL and DLGBLUP models improved prediction accuracy, when compared to GBLUP. The possibility of nonlinear relationships between traits offers a different perspective into multi-trait evaluations and prediction, as well as into the traits’ evolution over generations, with potential to further improve selection strategies in commercial livestock breeding programs. Moreover, DLGBLUP shows that DL can be used as a complement to statistical methods, by enhancing their performance.

## Background

Genomic prediction (GP) [1] in genetic evaluations uses DNA marker information, most commonly single nucleotide polymorphisms (SNPs) data, to predict the genetic merit of complex traits, referred to as genetic value (GV). When focusing on additive genetic values transmitted to the next generation, they are commonly termed as breeding values (BVs) in both livestock and plant studies. Traditionally, statistical methods applied to GP rely on linear mixed models [2] using either the pedigree information or SNP-genotypes, or both joined into the single-step method [3]. The predicted genetic values (PGV) are obtained under the assumption of normally distributed effects, either through the best linear unbiased prediction (BLUP), or Bayesian approaches, with their various prior assumptions and alphabets [1, 4]. These statistical methods, combined with the current availability of genomic information are very powerful, and have undoubtedly revolutionized genetic evaluations. Due to the assumption of normality of the data, when extended to a multi trait (MT) scenario, the relationship between traits is assumed to be linear [5, 6, 7]. If traits present nonlinear genetic relationships, the prediction accuracy of the current statistical methods is limited, despite the efforts of improvement by providing highly informative data, such as denser SNP-chips, or using a single-step strategy that combines the pedigree to the genomic data into a blended model [3].

The restriction to linearity, in addition to the continuously increasing amount of recorded and genotyped animals, which posed computational constraints for genetic evaluations with the classical statistical models, were among the reasons that pushed geneticists to explore the use of deep learning (DL) [8] for genetic evaluations and genomic predictions [9]. DL approaches [8] use artificial neural networks, which have the capacity to learn complex patterns and features from very large datasets, in order to map input to output data. Depending on the prediction task to achieve, different DL architectures have been considered, see e.g. [10, 11, 12, 13, 14].

Most studies that use DL for GP have focused on the inclusion of non-additive genetic effects (epistasis and dominance) with the objective of increasing prediction accuracy by accounting for such effects [15, 16, 17, 18, 19, 20, 21, 22, 23, 24]. More recently, Lee et al. [25] proposed deepGBLUP, a combination of DL and GBLUP into a single model. Given input SNP data, the DL networks of deepGBLUP extract the effects of adjacent SNPs using locally-connected layers to estimate a GV through fully-connected layers. In parallel, a GBLUP model accounting for additive, dominance and epistasis effects (through their respective genetic relationship matrices) is fitted. The final estimated GV is the sum of all previous estimated GV (DL, additive, dominance and epistasis). Applied to a real dataset of Korean native cattle, and to simulated data, the results of the proposed deepGBLUP model outperformed those of the traditional GBLUP and Bayesian methods in different single trait prediction scenarios.

In livestock production, beyond predicting GV for individual traits, breeders aim to jointly improve multiple traits of commercial interest, in order to achieve genetic progress for these traits altogether. When working with multiple traits, genetic correlations between traits [26] must be considered, since selection for one trait will affect the other correlated traits [27]. Genetic correlations are thus a relevant factor to account for, on the different stages of a genetic evaluation, from the estimation of variance components to the prediction of GV, in order to optimally improve multiple traits of interest altogether. It has been shown, for example, that it is more advantageous to perform a multi-trait model for genetic evaluation, that explicitly considers the genetic relationships between traits to improve their predictive ability [28].

Although non-linear correlations are not straightforward to handle by statistical methods, such hypothesis is suitable for the implementation of DL methods. Few studies explored the possibility of non-linear genetic correlations between traits [29, 30], and even so, such studies focused merely on how to identify these non-linear genetic correlations, rather than on how to account for them in GP.

The aim of this study was to explore the impact of non-linear genetic correlations between traits on the performance of GBLUP and DL methods. Using simulated data, we explored (1) how the presence of inter-trait non-linear correlations affect the performance of GBLUP with respect to the accuracy of the PGV, (2) how to use DL to model these non-linear correlations, and (3) how the presence of these correlations can affect the selection and the genetic progress over generations when ignored (GBLUP) or when they are taken into account (DL methods). We proposed a hybrid DL model, called DLGBLUP, that combines both DL and GBLUP, thus benefiting from their strengths while minimizing their pitfalls. DLGBLUP consists of two steps: first the GVs are predicted using a multi-trait GBLUP based on the genomic data, then GVs are re-predicted using a DL model that capture potential non-linear genetic correlations between traits.

## Methods

### Simulated data

The complete simulated genomic data consisted of 10,000 SNPs distributed across 29 chromosomes, with an average LD pattern resembling that of a cattle population. From the 10,000 simulated SNPs, 512 were assigned as quantitative trait loci (QTL) to be shared between all traits. The non-centered SNPs were coded as 0, 1, and 2, referring to homozygous for reference alleles, heterozygous, and homozygous for alternate alleles, respectively.

A quantitative reference trait was simulated using the 512 QTL for a total of 25,000 individuals, following the model: *y*_*ref*_ = *M* α_*ref*_ + *e*_*ref*_ = *g*_*ref*_ + *e*_*ref*_, where *M* is a matrix of which element M_i,j_ corresponds to the centered genotype of individual i at QTL j, α_*ref*_ = [α_ref,1_ …. α_ref,q_] is the vector of the q=512 additive QTL effects, such that α_ref,j_∼ N(0, σ^2^_αref_) i.i.d. for every j = 1, …, q, *g*_*ref*_ = *M*α_*ref*_ is the vector of the true genetic values (TGV) and *e*_*ref*_ ∼ N(0, σ^2^ I) is the vector of random errors. The genotypes and the reference trait were simulated using the GenEval R package [31].

Five dependent traits, were simulated conditional to the TGV of the reference trait as *y*_*t*_ = *g*_*t*_ + *e*_t_, for every t = 1, …, 5, such that *g*_*t*_= f_t_(*g*_*ref*_) + *g*_*t*_2_, where f_t_ is the function describing the (potentially) nonlinear relationship between the TGV of two traits. *g*_*t*_2_ is the vector of the genetic value specific to each dependent trait, simulated as *g*_*t*_2_ = *M*_t_α_t_, where *M*_*t*_ is the genotype matrix of QTLs specific to trait t and different from the common QTLs and different between the dependent traits; α_*t*_∼ N(0, σ^2^ I) is the vector of the corresponding additive QTL effects. *g*_*t*_ was normalized such as σ^2^_gt_ = σ^2^_yt_ × h^2^, where h^2^ is the heritability. The error vector *e* ∼ N(0, σ^2^_et_ I). We fixed σ^2^_y_ = 20 for all traits. Different values of h^2^(0.6, 0.3, 0.15, 0.05) and different numbers of specific QTLs (0, 10, 50, 250) were considered for the traits. While different levels of h^2^ and number of specific QTL were tested, the values were maintained the same between traits of each set. Four relationships were considered for the dependent traits: (1) linear, (2) quadratic, (3) logistic and (4) sinusoidal. To assess the repeatability of our findings, we simulated 20 replicas of the complete data set under each combination of different levels of h^2^ and number of QTL.

To train and evaluate the DL models, the dataset was split into three sets: training (80%), validation (10%), and test set (10%), being the test set that for which PGV are to be obtained in the absence of phenotypic records, while both the training and validation sets are those for which individuals have genotypes and phenotypic records. Different from statistical models, DL requires an internal validation set with the complete information to fine-tune the model parameters. For the GBLUP model, the training and validation sets were combined to fit the model, and the same test set (10% of individuals) was kept for evaluation.

### Real Data

The real dataset consisted of 113,599 genotyped Holstein cows, with a total of 53,469 SNPs post-editing (original SNP data from the EuroGMD v1, a customized ILLUMINA genotyping microarray that contains approximately 70,000 SNPs). For the performance records, we utilized the yield deviations (YD) of 33 traits: five dairy production traits (heritability 38–63%), somatic cell score (heritability 19%), six fertility traits (heritability 1–7%), and 21 dairy morphology traits (heritability 10–51%). The YD consisted of the corrected phenotype for every non-genetic effect from single-trait models used in the French genetic evaluation (effects were adequately considered according to each one of the traits) and were obtained using the single-step GBLUP (ssGBLUP). By considering the generation between animals, we split the dataset in three sets: training (82%), validation (9%), and test set (9%). A cow from any of the three sets has no paternal half-sisters in the other two sets. Cows in test set are more recent than those in validation set, and cows in validation set are more recent than those in training set.

### Multi-Trait Genomic Prediction

We used three models to perform GP: a baseline GBLUP, a DL model, and a hybrid DLGBLUP model that combines the two previous ones.

### Genomic Best Linear Unbiased Prediction (GBLUP)

The GBLUP model is one of the most popular statistical methods used to predict the GV of genotyped individuals using their genomic relationship matrix. The model considered for multi-trait GBLUP is:

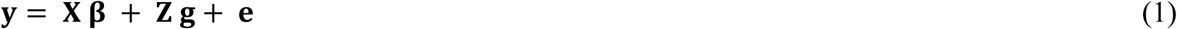

where Y = [Y_1_ … Y_T_] is a matrix of the phenotype vectors for all trait included in the model; X is a design matrix for the fixed effects, β = [β_1_ … β_T_], such that β_t_ is the vector of fixed effects for trait k = 1, …, K; Z is an incidence matrix linking the phenotypic records to the genetic values *g* = [*g*′_1_ … *g*′_T_]′, such that *g*_t_ is the vector of genetic values for trait *t*, with distribution,

*g*∼ N(O, Σ_g_⊗G) in which Σ_g_ is the additive genetic (co)variance matrix of the traits, G is the genomic relationship matrix (GRM) [2], and ⊗ represents their Kronecker product; and *e* = [*e*′_1_ … *e*′_T_]′ is the matrix of normally distributed random errors with *e* ∼ N(O, Σ⊗I), in which **R** is the error (co)variance matrix between the traits. We used the R (4.1.1) package BGLR package (1.0.8) [32] to perform the multi-trait GBLUP on the simulated data, here we did not consider a fixed effect so β is just the vector of the intercepts. For the real dataset, a single-trait ssGBLUP analysis was performed using the HSSGBLUP software [33], developed for the French cattle genetic evaluation.

### Deep Learning model

The Multi-Layer Perceptron (MLP) [34] is a feedforward network that maps an input to an output using learned parameters. The model transforms the input by passing through multiple connected layers (hidden layers) up to the output. In a general execution, each node is characterized by the following equation:

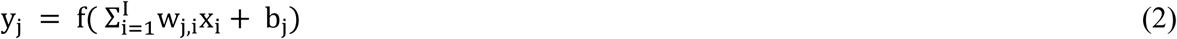

where y_j_ is the scalar output of node j, b_j_ is the bias, x_i_ represents the output of node i ∈ {1, …, I} of the previous layer, w_j,i_ is the weight assigned to this x_i_, and f is a nonlinear activation function.

During training, the optimization process adjusts the weights and biases of the MLP by minimizing a loss function that quantifies the difference between the predicted and true outputs on the training set. The Mean Squared Error (MSE) and Mean Absolute Error (MAE) losses are commonly used for regression, while cross entropy loss is used for classification. In practice, stochastic gradient descent is used to update the parameters (weights and biases).

The proposed DL model, illustrated in Figure 1, comprises two distinct modules: SNPs2Trait and Trait2Trait. The first module, SNPs2Trait, takes the SNP data as input, and provides a first predicted GV (PGV__1_) for multiple traits as an output by capturing the relationship between the genomic data and the phenotype. The hidden layers enable the construction of a latent genome representation common to all traits. This latent representation is used in the output layer to predict each trait separately. The second module Trait2Trait takes as input the PGV__1_ of all traits and outputs a new predicted GV (PGV__2_) for all traits by capturing the relationship between all traits that will be considered as the final prediction of the model.

**Figure 1:**
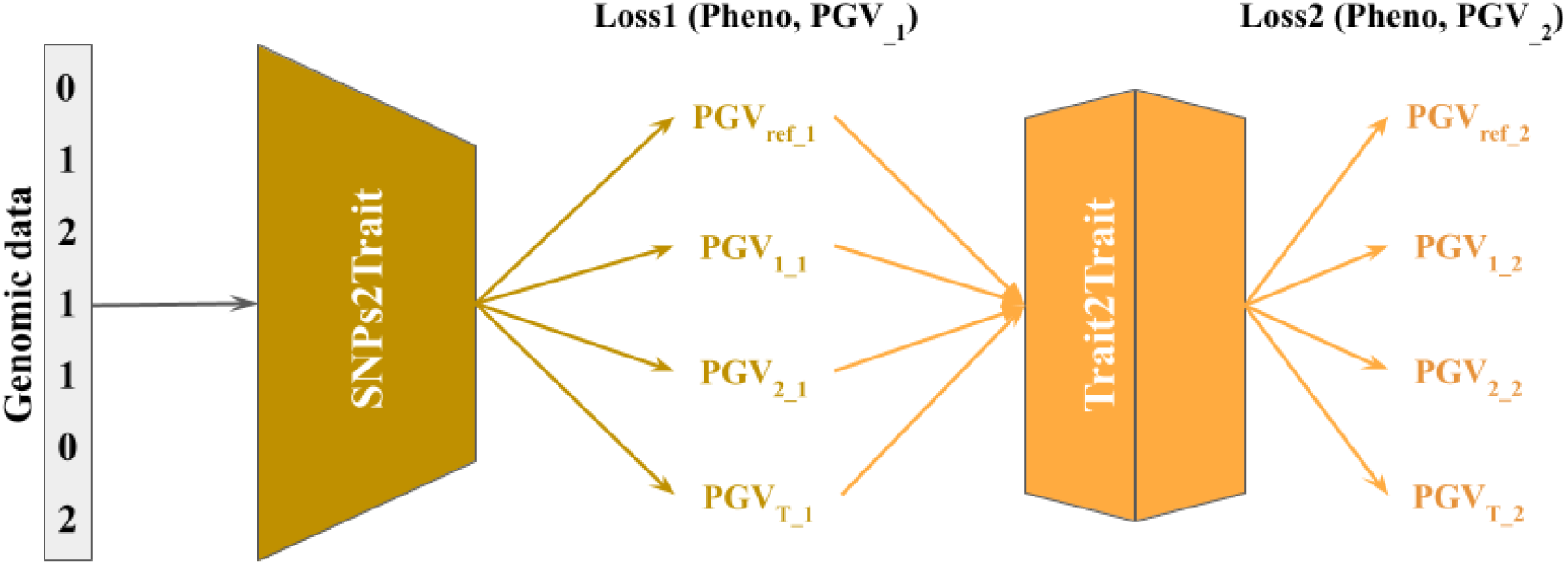
Deep Learning Genomic Prediction Frameworks.

The SNPs2Trait module consists of one hidden layer (with 400 neurons) and the Trait2Trait module consists of two hidden linear layers (with 400 and 256 neurons, respectively). LeakyReLU activation function was used with a fixed negative slope of 0.1. The model was trained for 100 maximum epochs, with each epoch consisting of a single pass through the training dataset, using a batch size of 200. An early stopping was employed to terminate training if there was no improvement in validation loss after 10 epochs. As the loss function, we used Huber loss [35] due to its combination of the advantages of both MSE and MAE, providing a balance between sensitivity to small errors and robustness to outliers. This loss is described as:

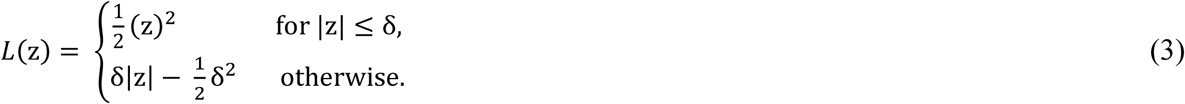

where z is the error and δ (fixed to 1) is a threshold parameter that dictates the transition between the two losses. We used Adam optimizer (Adaptive Moment Estimation) with a learning rate of 10^−4^. The choice of optimizer, learning rate, activation function and number of neurons in hidden layers was made after a grid search (Table S1).

Two training approaches were used: (1) training the two modules as one model so the loss function was the sum of loss from each module, i.e.,

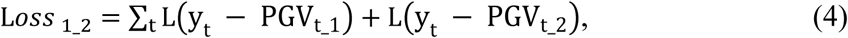

where y is the true phenotype value for trait t; (2) training modules sequentially in terms of gradient flow: the first module was trained initially, and once this was completed, its parameters were fixed by stopping the gradient flow. This procedure isolated it from the training phase of the second module, maintaining the integrity of the initial module’s parameters as the later module was trained. The model was implemented using pytorch (1.10.2) in python and trained on a single GPU with 48 GB memory (NVIDIA A40).

### DLGBLUP model

We proposed a hybrid version of the previous model called DLGBLUP (Figure 2), where the first module is replaced with GBLUP. The reason behind this variation is that the first module, SNPs2Trait, performs a similar function to GBLUP by learning the patterns between genotype and phenotype to predict the genetic value (GV). We wanted to test the effect of starting with the results of GBLUP to make the final predictions using the Trait2Trait module, which remains unchanged.

**Figure 2:**
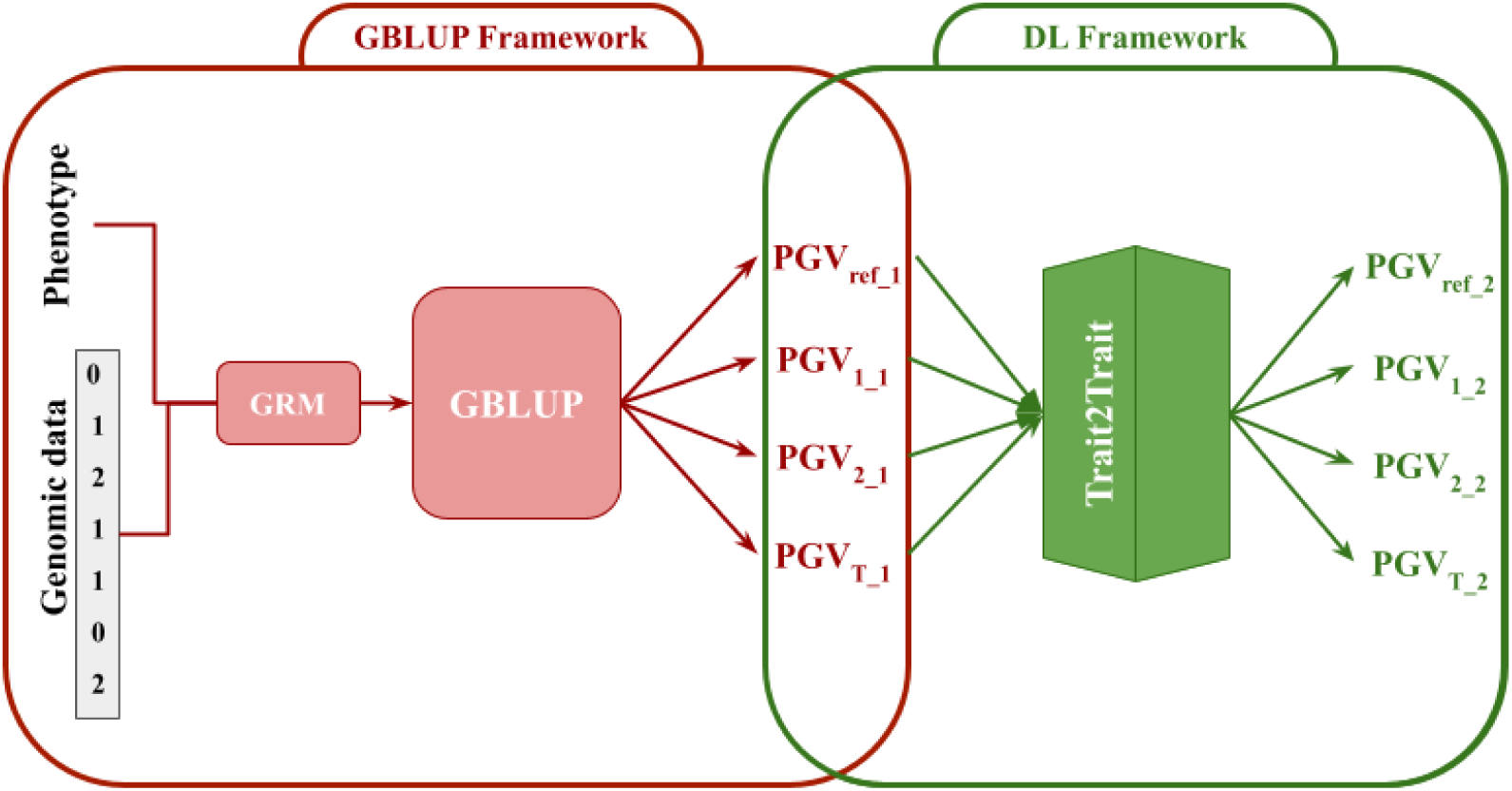
DLGBLUP Genomic Prediction Frameworks.

The GBLUP in red and DL in green.

### Inter-trait relationship representation

In order to represent the relationship between the reference trait and a dependent trait, a MeanTrait2Trait module was proposed (Figure 3). This module outputs predictions of the dependent traits which we call PGV__3_ by taking PGV of reference trait (PGV _ref_) as input. As such, MeanTrait2Trait can capture the mean relationship between reference trait (input) and the other dependent traits. The training of MeanTrait2Trait was done using *Loss*_3_ = ∑_t_ L(PGV_t_ − PGV_t_3_). This last module is used for illustration purpose only.

**Figure 3:**
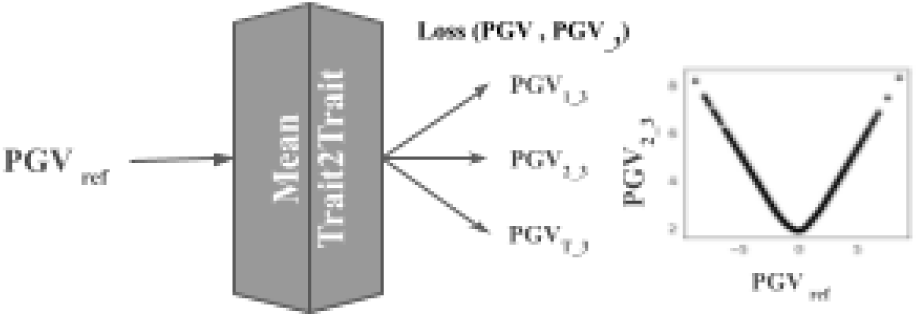
Mean Trait2Trait model

### Impact of QTL Exclusion and Non-Causative SNP Addition

We first considered the ideal case where only the true QTL were provided to the prediction methods which represent our baseline. To evaluate the impact of including SNPs that were not QTL on the prediction accuracy, we gradually added non-causative SNPs to the input genomic data. This resulted in scenarios where the genomic composition included 10%, 25%, 50%, and 75% non-QTL SNPs. Finally, a final scenario consisted of using only non-causative SNPs, excluding all the QTL. A comparison of the results from a model excluding non-causative mutations, but including only half of the QTL was also included in this study. These scenarios offer a clear insight into how the signal-to-noise ratio impacts the prediction of PGV and the effect of relationships between traits.

### Genomic Selection

After performing the multi-trait genomic prediction, we computed a selection index (SI) as:

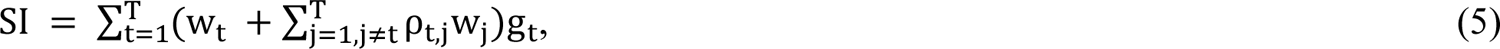

in which w_t_’s are the weights given to traits t = 1, …, T, ρ_t,j_ is the empirical correlation between the GVs of traits t and j, and g_t_’s are the GVs of trait t = 1, …, T. For this study, we assigned equal weights to all traits in order to compare the general evolution of the genetic gain based on the breeding values obtained with the different models. Based on the SI, at each generation the top 10% eligible males were selected and mated with all females from the latest generations, in order to generate the next generation, maintaining at each new generation a female-to-male proportion of 80%-20%. Males were kept for breeding across four generations, while females were kept for two generations only.

This selection and mating design were chosen to mimic the breeding program of a dairy cattle population. A base population of independent individuals (i.e. generation zero, or G0) was considered to start the selection process, followed by seven simulated generations under selection referred to as G1, …, G7.

This simulation scheme under selection allowed us to both study the evolution of the nonlinear genetic relationships between traits when the population is under selection, and compare the genetic gain of selection, based on the additive GV obtained with the different models, using their values to construct the SI in equation (5). Because of the way we simulate the data, the PGV obtained by all prediction models could express non-additive effects. Since we are interested in the additive PGV, we computed it as follow: at each generation we simulated the genotype of 20 offspring for each eligible male based on random mating schemes with the females. Then, the PGVs of these simulated offspring were predicted with the different models (GBLUP, DL, and DLGBLUP) trained on a dataset where all individuals until the latest generation were genotyped and phenotyped. The additive PGV of the eligible males were finally computed as the average of the PGVs of their simulated offspring. The GVs for the reference trait on all subsequent generations were simulated using the same original QTL effects of G0, and the other dependent traits were simulated as previously described in the ‘Simulated Data’ section. The number of individuals in each of the generations G1 to G7 was maintained as 5,000. With each new generation, the reference population to perform the prediction models was updated to comprise all individuals from G0 to the latest with both genotypes and phenotypic records, and then used to perform the genetic evaluation. For DL and DLGBLUP, the genetic evaluation model with a new generation was initialized using the weights (DL parameters) obtained at the evaluation of the previous generation.

### Evaluation metrics

To evaluate the performance of each model in predicting genetic values, we used the Pearson correlation coefficient to calculate the prediction accuracy for each individual trait, as well as the MSE between the TGV and PGV, and performed a visual assessment of the relationship form between traits. We also computed the contribution percentage of the variation in linear and nonlinear relationships to the variation in the predicted genetic value of dependent traits. This was done by calculating the variance of the linear or nonlinear predictions using the MeanTrait2Trait model of the dependent trait and dividing it by the mean of the predicted genetic value of the same trait multiplied by 100. To evaluate the performance of each model with respect to genomic selection, we used the genetic gain computed as the difference between the mean of the TGV for individuals in G0 and the mean of the TGV for individuals in the next generation.

## Results

Here we present the results of DL-based model with MLP architecture and trained following the second training approach (sequential training) which gave better results. For the simplicity, we called the traits based on the form of relationship they have with the reference trait, for example quadratic trait correspond to the trait with f as quadratic. We used the phenotype set with heritability 0.3 and 50 specific QTLs as default. The results of sinusoidal trait can be found in Figure S2 as it was used to test an extreme case of non-linearity.

### Genomic Prediction

#### Prediction Accuracy

Figure 4 presents the boxplots comparing the performance of the GBLUP, DL, and DLGBLUP models using only the QTL as input data, with respect to the prediction accuracy and the MSE, over the 50 replicates performed.

**Figure 4:**
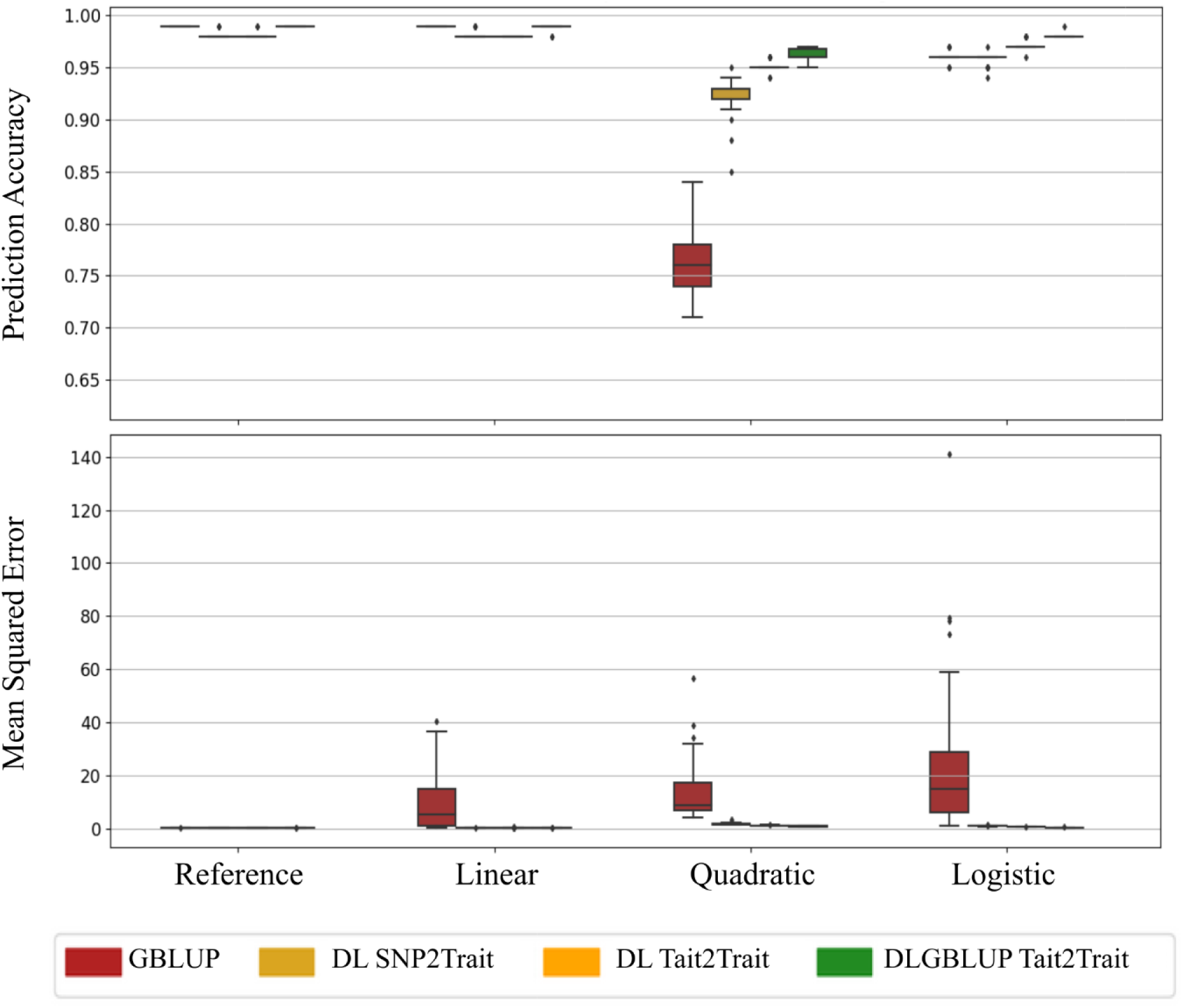
Performance comparison of different prediction models for various traits. GBLUP (red), Deep Learning (DL) (yellow); with the two modules SNPs2Trait and Trait2Trait and DLGBLUP (green). The prediction was made using genomic data with just QTL.

At the additive effect level, GBLUP demonstrated superior prediction accuracy compared to the DL SNP2Trait model for reference and linear trait. The opposite was shown for the quadratic trait. For logistic trait all methods exhibited similar results. Nonetheless, DL SNPs2Trait presented a consistently lower MSE for all traits except the reference trait as expected due to the shrinkage of the marker effects in the GBLUP procedure.

Independent of the procedure used to obtain PGV_1 (either GBLUP or DL SNP2Trait), the application of theTrait2Trait module always improved the prediction accuracy of PGV_2 (Figure 4). The prediction accuracy increased by 0.01 to 0.07 for DL Trait2Trait module, and by 0.01 to 0.2, depending on the type of nonlinear correlation between traits. After passing through the Trait2Trait module, no further improvement was provided for the prediction accuracy of the linear traits (reference included), which were already well predicted specially by GBLUP.

DLGBLUP outperformed DL Trait2Trait by a margin of 0.01 to 0.02 in terms of prediction accuracy. Of all the evaluated models, DLGBLUP Trait2Trait consistently yielded the lowest MSE across all dependent traits, with more reliable and precise estimation, with a prediction accuracy equal or higher than all the other models.

### Inter-trait relationship representation

The MeanTrait2Trait module represent the relationship between traits correctly, whether it was linear or nonlinear, using the PGV _ref_ predicted from any prediction method. Moreover, the shape of the PGV was adjusted according to the trait’s relationship through the Trait2Trait part. The SNPs2Trait module was able to identify a slight non-linearity between the traits but not the complete shape detected by Trait2Trait. A more accurate input, i.e. the GBLUP predictions, to the Trait2Trait module, enabled the prediction of a clearer and more precise relationship between traits, as illustrated in Figure 5 and Figure S2. Additionally, Trait2Trait was able to detect the genetic relationships between traits, while GBLUP was able to do so only for the linear trait. For the other nonlinear traits, we observed that GBLUP transformed their relationship to be linear, sometimes identifying a level of linear relationship (logistic trait), or completely missing any relationship between traits (quadratic trait). When examining the contribution of this relationship to the total genetic value, we notice a high percentage of nonlinear contribution for nonlinear traits with a large difference with linear contribution, especially for quadratic traits. There is a significant contribution from the linear relationship for linear models predicting linear traits, which is higher than the nonlinear contribution (Table 1).

**Figure 5:**
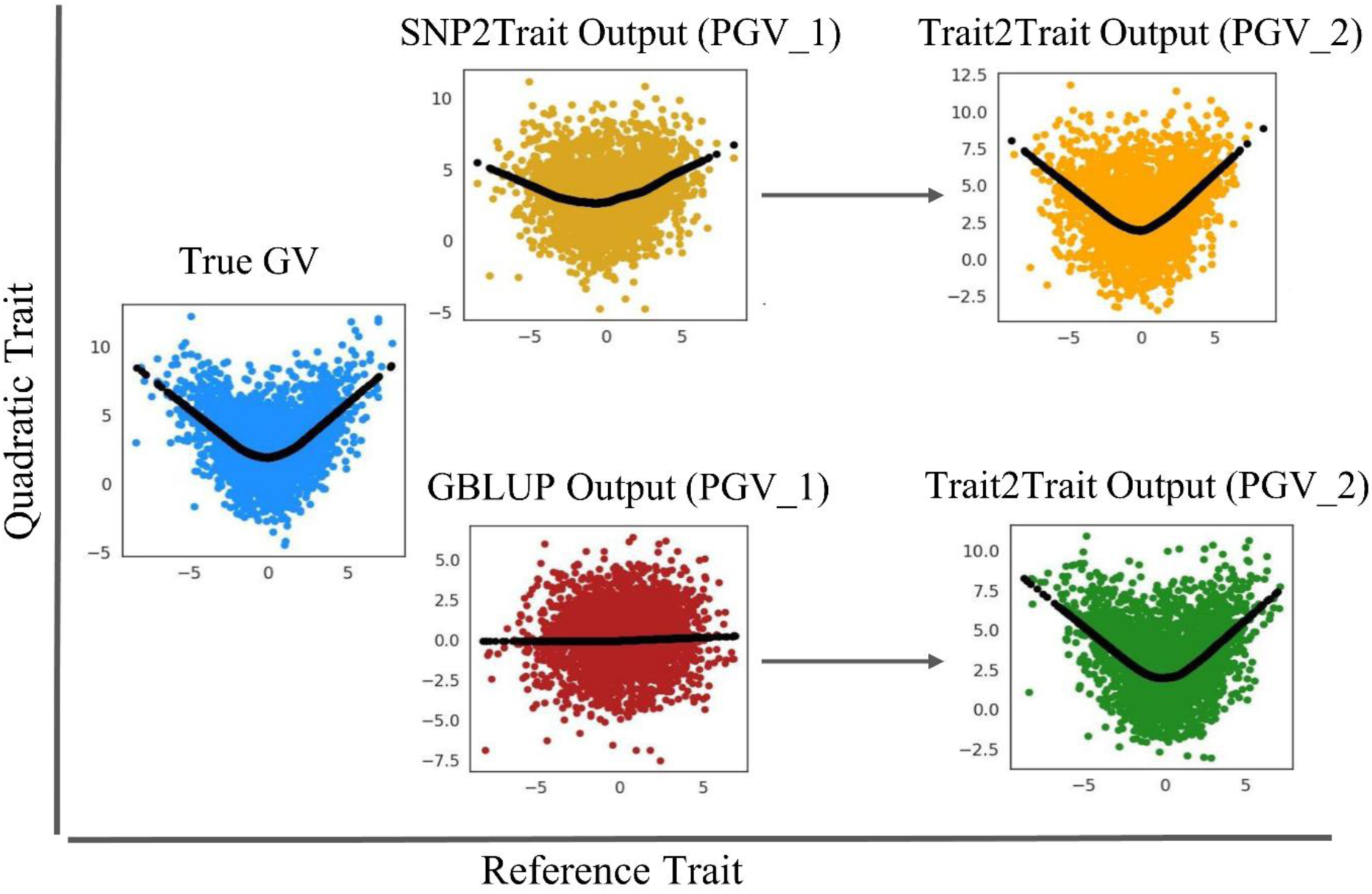
Comparison plots of predicted genetic values for quadratic trait with SNP2Trait and Trait2Trait modules of DL model and GBLUP and Trait2Trait modules of DLGBLUP. The True GV in blue, DL SNP2Trait prediction in light yellow, DL Trait2Trait prediction in orange, GBLUP prediction in red, and Trait2Trait DLGBLUP in green.

**Table 1:** Contribution of Linear and nonlinear relationship prediction for simulated data.

| Contribution % |  |  |
| --- | --- | --- |
| Trait | Linear | Non-linear |
| Linear | 24.4 | 21.58 |
| Quadratic | 0.26 | 34.68 |
| Logistic | 17.18 | 24.17 |

### Effect of Heritability, number of specific QTL, and level of Genetic Relationship

Heritability directly influences the quality of GV predictions from additive effects. The higher the heritability, the higher the prediction accuracy of GBLUP, as well as the prediction accuracy made by the Trait2Trait part and its improvement in comparison to GBLUP, for the nonlinear traits, reaching 0.17 for the quadratic traits with heritability of 0.6 (Table 2).

**Table 2:** Prediction accuracy comparison of different prediction models with different heritability with 50 specific QTL.

| Heritability | Model | Traits |  |  |  |
| --- | --- | --- | --- | --- | --- |
|  |  | Reference | Linear | Quadratic | Logistic |
| 0.05 | GBLUP | 0.83 | 0.8 | 0.8 | 0.84 |
|  | DL | 0.72 | 0.67 | 0.7 | 0.74 |
|  | DLGBLUP | 0.82 | 0.79 | 0.8 | 0.83 |
| 0.15 | GBLUP | 0.93 | 0.94 | 0.82 | 0.93 |
|  | DL | 0.88 | 0.89 | 0.81 | 0.89 |
|  | DLGBLUP | 0.93 | 0.93 | 0.87 | 0.92 |
| 0.3 | GBLUP | 0.97 | 0.97 | 0.84 | 0.96 |
|  | DL | 0.95 | 0.95 | 0.91 | 0.94 |
|  | DLGBLUP | 0.97 | 0.97 | 0.94 | 0.95 |
| 0.6 | GBLUP | 0.99 | 0.99 | 0.79 | 0.96 |
|  | DL | 0.98 | 0.98 | 0.96 | 0.97 |
|  | DLGBLUP | 0.99 | 0.99 | 0.96 | 0.98 |

Moreover, there was an effect of the number of specific QTL that control the genetic relationship (Figure 6). The higher the number of specific QTL the less clear the relationship between traits. A higher genetic relationship - a lower number of specific QTL in our case-intensifies the nonlinear effects, reducing GBLUP’s prediction accuracy while simultaneously increasing the potential for improvement via the Trait2Trait component, as shown in Table 3.

**Figure 6:**
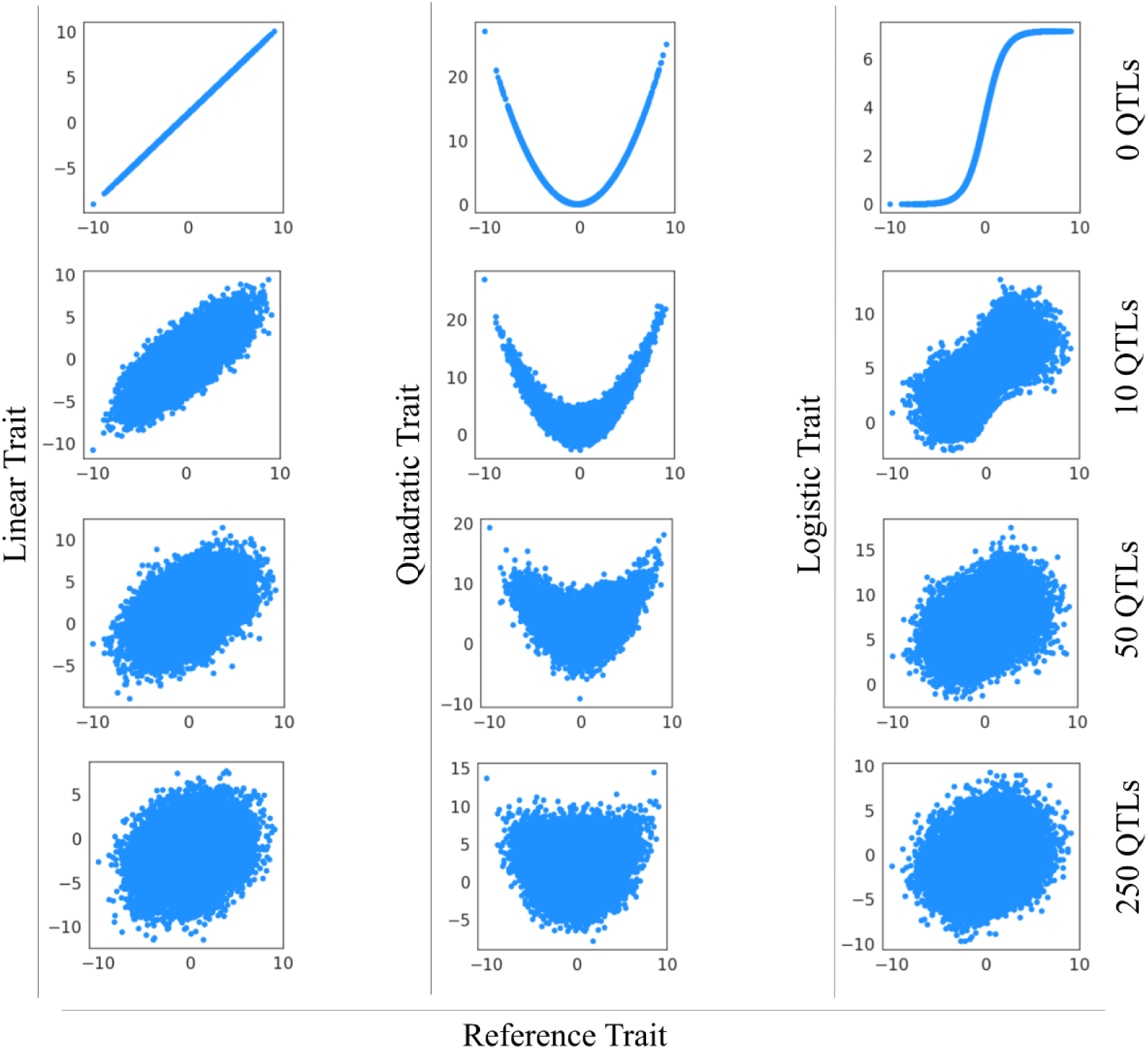
Comparison plots of true genetic relationship between traits with different number of specific QTLs. The reference trait is represented in x-axis and all dependent traits in y-axes. The blue points represent the True GV of a population.

**Table 3:** Prediction accuracy comparison of different prediction models with different number of specific QTL and heritability 0.3.

| QTLs | Model | Traits |  |  |  |
| --- | --- | --- | --- | --- | --- |
|  |  | Reference | Linear | Quadratic | Logistic |
| 0 | GBLUP | 0.99 | 0.99 | 0.13 | 0.94 |
|  | DL | 0.97 | 0.97 | 0.85 | 0.93 |
|  | DLGBLUP | 0.99 | 0.99 | 0.95 | 0.97 |
| 10 | GBLUP | 0.98 | 0.98 | 0.42 | 0.95 |
|  | DL | 0.96 | 0.96 | 0.82 | 0.94 |
|  | DLGBLUP | 0.98 | 0.97 | 0.9 | 0.96 |
| 50 | GBLUP | 0.97 | 0.97 | 0.84 | 0.96 |
|  | DL | 0.95 | 0.95 | 0.91 | 0.94 |
|  | DLGBLUP | 0.97 | 0.97 | 0.94 | 0.95 |
| 250 | GBLUP | 0.92 | 0.93 | 0.9 | 0.93 |
|  | DL | 0.89 | 0.89 | 0.86 | 0.89 |
|  | DLGBLUP | 0.92 | 0.92 | 0.91 | 0.93 |

### Impact of QTL Exclusion and Non-Causative SNP Addition

Table 4 presents the impact on the prediction accuracy of incorporating SNPs that were not QTL in the genomic data used. The most accurate predictions were achieved when the input data comprised exclusively all the QTLs implicated in the TGV of a trait, as expected, with DLGBLUP achieving the best predictions. When excluding half of the QTLs, or when adding non-causative SNPs to the input genotypes the prediction accuracy decreased. The decrease in prediction accuracy when non-causative SNPs were introduced was progressive as more and more SNPs were included to the genomic data, which results in a decrease in the advantage of DLGBLUP. Excluding all QTL, and keeping only non-causative SNPs in LD with the QTL, resulted in PGV with accuracies ranging from 0.55 to 0.7, and an improvement made by DLGBLUP ranging from 0.01 to 0.03, when compared to both DL and GBLUP.

**Table 4:**
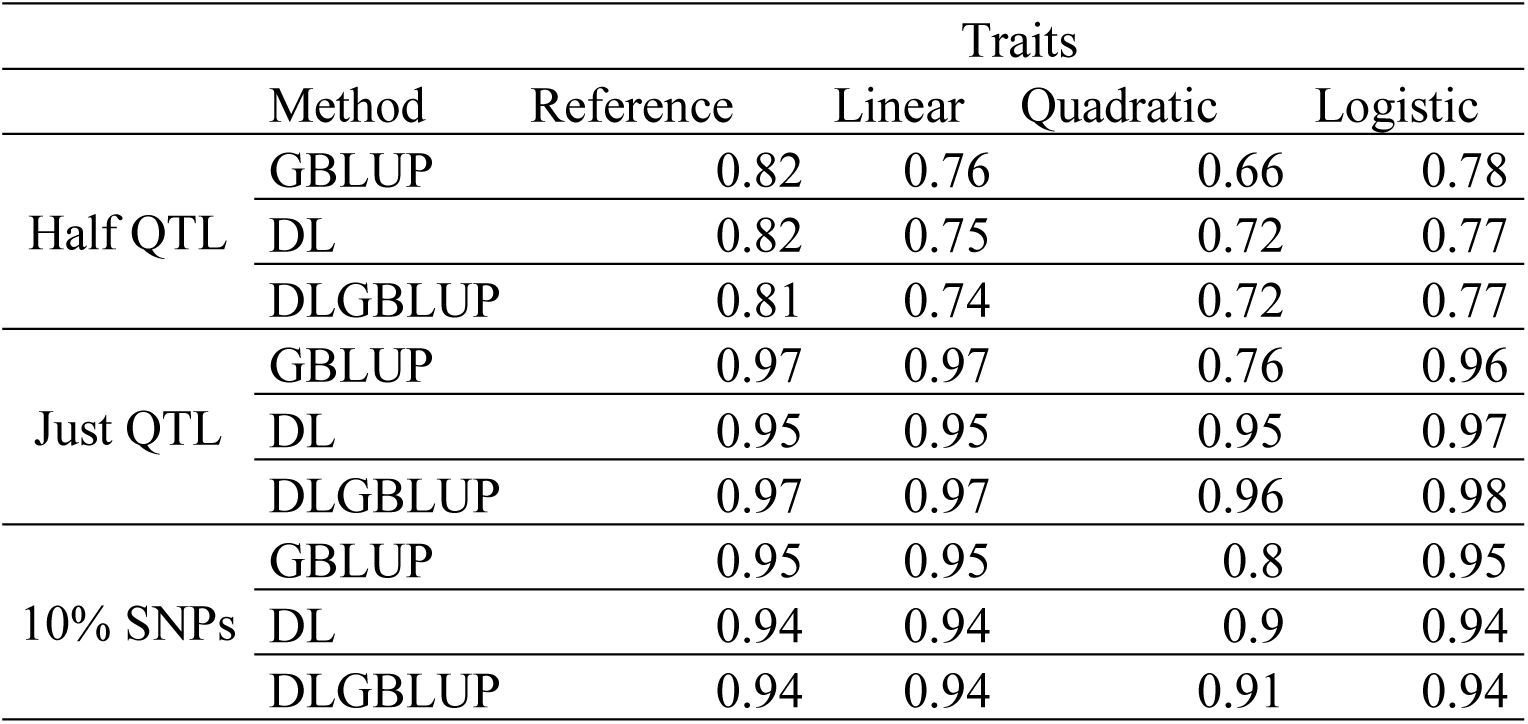
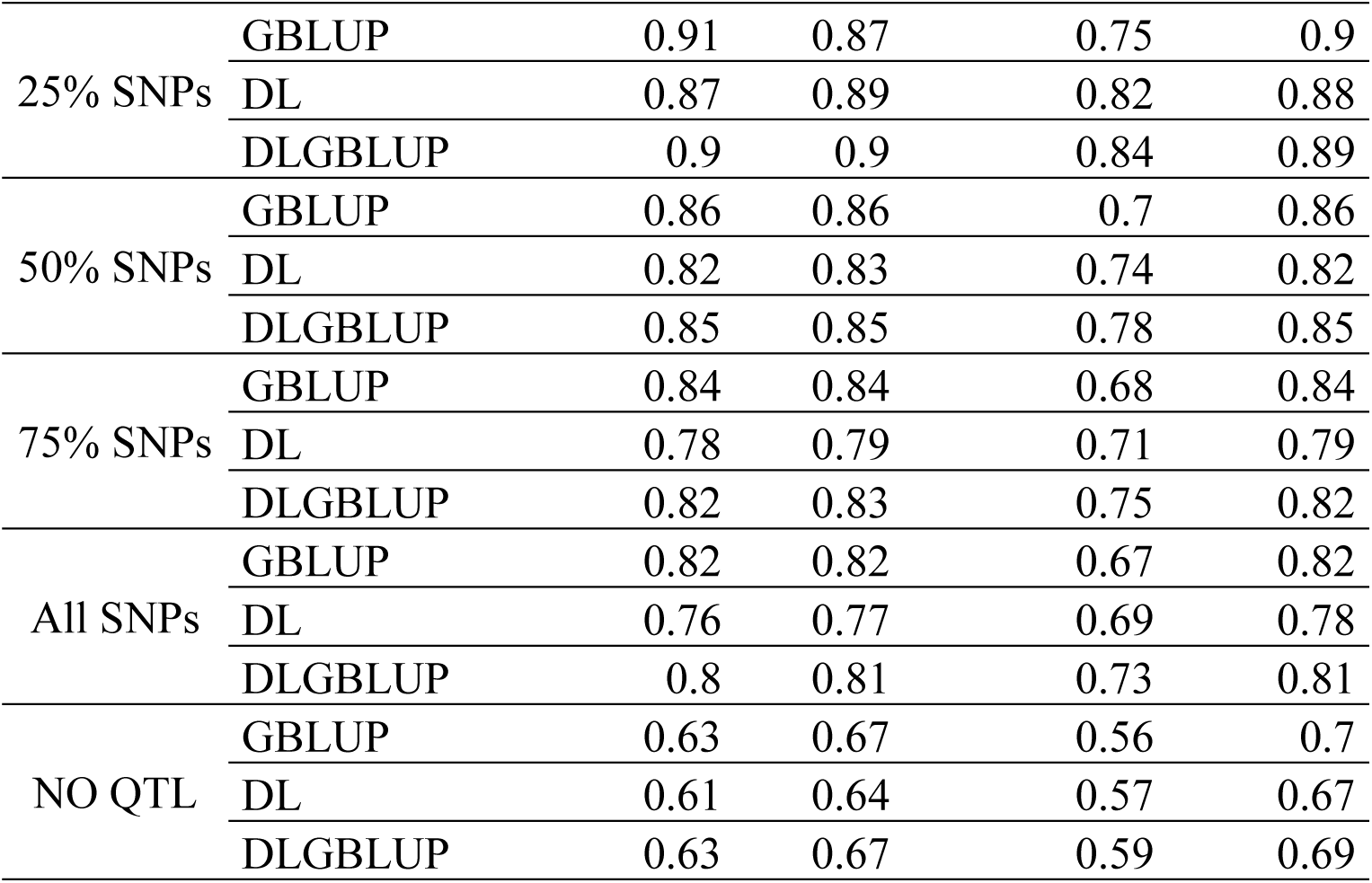
Prediction accuracy comparison of different prediction models using genomic data with different numbers of SNPs, a h^2^of 0.3 and 50 specific QTLs.

|  |  | Traits |  |  |  |
| --- | --- | --- | --- | --- | --- |
|  | Method | Reference | Linear | Quadratic | Logistic |
| Half QTL | GBLUP | 0.82 | 0.76 | 0.66 | 0.78 |
|  | DL | 0.82 | 0.75 | 0.72 | 0.77 |
|  | DLGBLUP | 0.81 | 0.74 | 0.72 | 0.77 |
| Just QTL | GBLUP | 0.97 | 0.97 | 0.76 | 0.96 |
|  | DL | 0.95 | 0.95 | 0.95 | 0.97 |
|  | DLGBLUP | 0.97 | 0.97 | 0.96 | 0.98 |
| 10% SNPs | GBLUP | 0.95 | 0.95 | 0.8 | 0.95 |
|  | DL | 0.94 | 0.94 | 0.9 | 0.94 |
|  | DLGBLUP | 0.94 | 0.94 | 0.91 | 0.94 |
| 25% SNPs | GBLUP | 0.91 | 0.87 | 0.75 | 0.9 |
|  | DL | 0.87 | 0.89 | 0.82 | 0.88 |
|  | DLGBLUP | 0.9 | 0.9 | 0.84 | 0.89 |
| 50% SNPs | GBLUP | 0.86 | 0.86 | 0.7 | 0.86 |
|  | DL | 0.82 | 0.83 | 0.74 | 0.82 |
|  | DLGBLUP | 0.85 | 0.85 | 0.78 | 0.85 |
| 75% SNPs | GBLUP | 0.84 | 0.84 | 0.68 | 0.84 |
|  | DL | 0.78 | 0.79 | 0.71 | 0.79 |
|  | DLGBLUP | 0.82 | 0.83 | 0.75 | 0.82 |
| All SNPs | GBLUP | 0.82 | 0.82 | 0.67 | 0.82 |
|  | DL | 0.76 | 0.77 | 0.69 | 0.78 |
|  | DLGBLUP | 0.8 | 0.81 | 0.73 | 0.81 |
| NO QTL | GBLUP | 0.63 | 0.67 | 0.56 | 0.7 |
|  | DL | 0.61 | 0.64 | 0.57 | 0.67 |
|  | DLGBLUP | 0.63 | 0.67 | 0.59 | 0.69 |

### Genomic Selection

#### Genetic relationship across generations

The prediction accuracy of the predicted additive genetic effects for all methods was similar, ranging between 0.9 and 0.97, as shown in Table 5. Figure 7.A shows the evolution of the genetic relationships between the TGV of the traits over the seven generations, under a multi-trait selection using the selection index (SI) based on the additive GV obtained with pure DL using only the QTLs as input data. As expected, the relationship between the reference trait and the linear trait remained linear over all the generations under selection. In contrast, the relationship between the reference trait and the quadratic trait evolved from nonlinear to linear very quickly, with its nonlinearity being almost imperceptible from G1. The logistic trait remained nonlinear, although, the shape of its relationship with the reference trait changed completely with respect to the original one in G0. These evolutions of the shape of the relationships persisted whether selection was based on any of the predicted additive GV (Figure S3).

**Figure 7:**
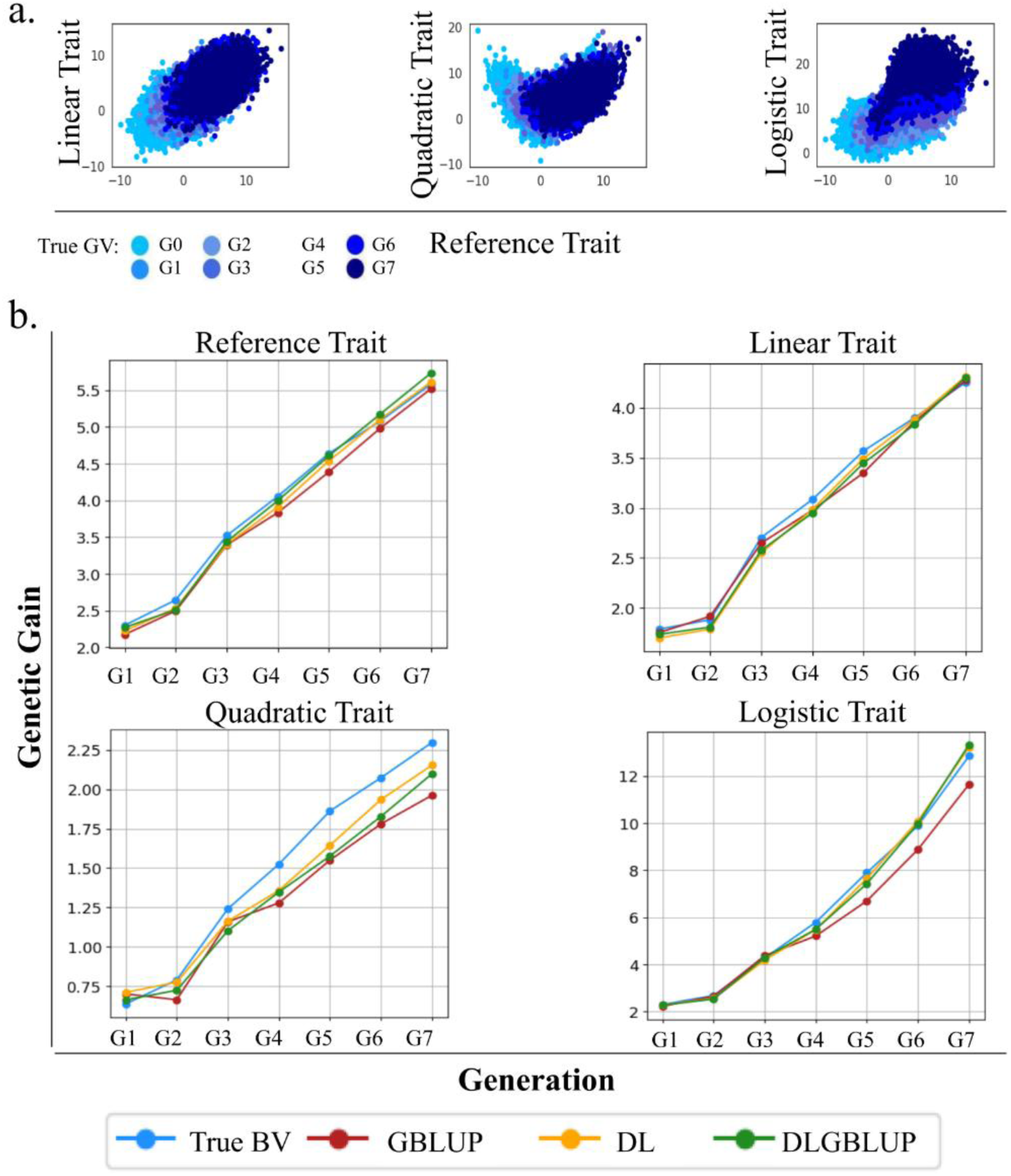
Comparative analysis of traits progression across generations. (A) Plots of relationship between the TGV of reference trait (x-axis) and dependent traits (y-axis) over 8 generations based on DL predictions. (B) Comparison between the Genetic Gain of 7 generations with 15% male selection using the true and the predicted additive GV with GBLUP, DL, and DLGBLUP for all traits.

**Table 5:**
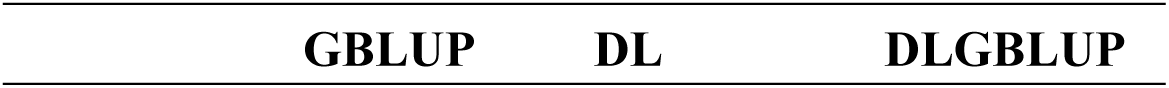
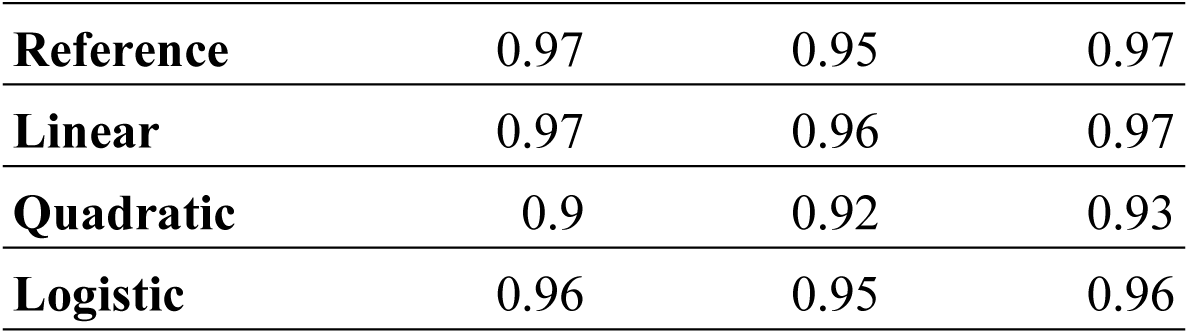
Prediction accuracy of additive genetic value comparison of different prediction models.

|  |  |  |  |
| --- | --- | --- | --- |
| **Reference** | 0.97 | 0.95 | 0.97 |
| **Linear** | 0.97 | 0.96 | 0.97 |
| **Quadratic** | 0.9 | 0.92 | 0.93 |
| **Logistic** | 0.96 | 0.95 | 0.96 |

#### Genetic Gain

Genetic gain was achieved for all the traits, following the selection of the top 10% males (Figure 7.B), based on the SI with any of the additive PGV. The magnitude of this gain varied depending on the form of relationship, with the most substantial gains observed for the logistic trait. For both reference and linear traits, the progress based on the SI using the additive GV from all models were equivalent. Conversely, for each of the nonlinear traits, the genetic progress achieved based on the SI using the additive GV from either DL or DLGBLUP, or both for some traits, was greater than the genetic progress achieved using the additive GV from GBLUP.

#### Application on Real data

DLGBLUP was applied on real data. The YD was used instead of phenotype in simulated data and the results of unitrait GBLUP of 33 traits was used as input for Trai2Trait module. Here we are representing the results of six traits: milk yield, fat yield, protein content as production traits, somatic cell score as health trait, conception rate cows as fertility trait, and depth of furrow as morphology trait (see additional files for other traits). On test set, no improvement was observed using Trait2Trait (Table 6 and Table S2).

**Table 6:** Prediction accuracy comparison between GBLUP and Trait2Trait module of DLGBLUP model for multiple traits for real data on test set.

| Trait | Heritability | GBLUP | Trait2Trait |
| --- | --- | --- | --- |
| **Milk Yield** | 0.38 | 0.65 | 0.65 |
| **Fat Yield** | 0.38 | 0.62 | 0.62 |
| **Protein Content** | 0.63 | 0.89 | 0.89 |
| **Somatic cell score** | 0.19 | 0.34 | 0.35 |
| **Conception Rate Cows** | 0.04 | 0.12 | 0.12 |
| **Depth of Furrow** | 0.23 | 0.38 | 0.38 |

Even if the relationship seems to be linear when plotted over the predicted genetic value on population level, MeanTrait2Trait was able of detecting a form of nonlinearity between different traits (Figure 8). For example, the relationship between milk yield and fat yield demonstrates a positive, linear-like trend, whereas the relationships between the other traits exhibit a more nonlinear pattern. However, when considering the contribution percentage, both linear and nonlinear relationships contribute only a small percentage for most traits, except for milk yield, which has approximately 6%. The contribution of nonlinearity was higher than that of linearity for all traits, except for the fertility trait, where the difference was 0.18% (Table 7).

**Figure 8:**
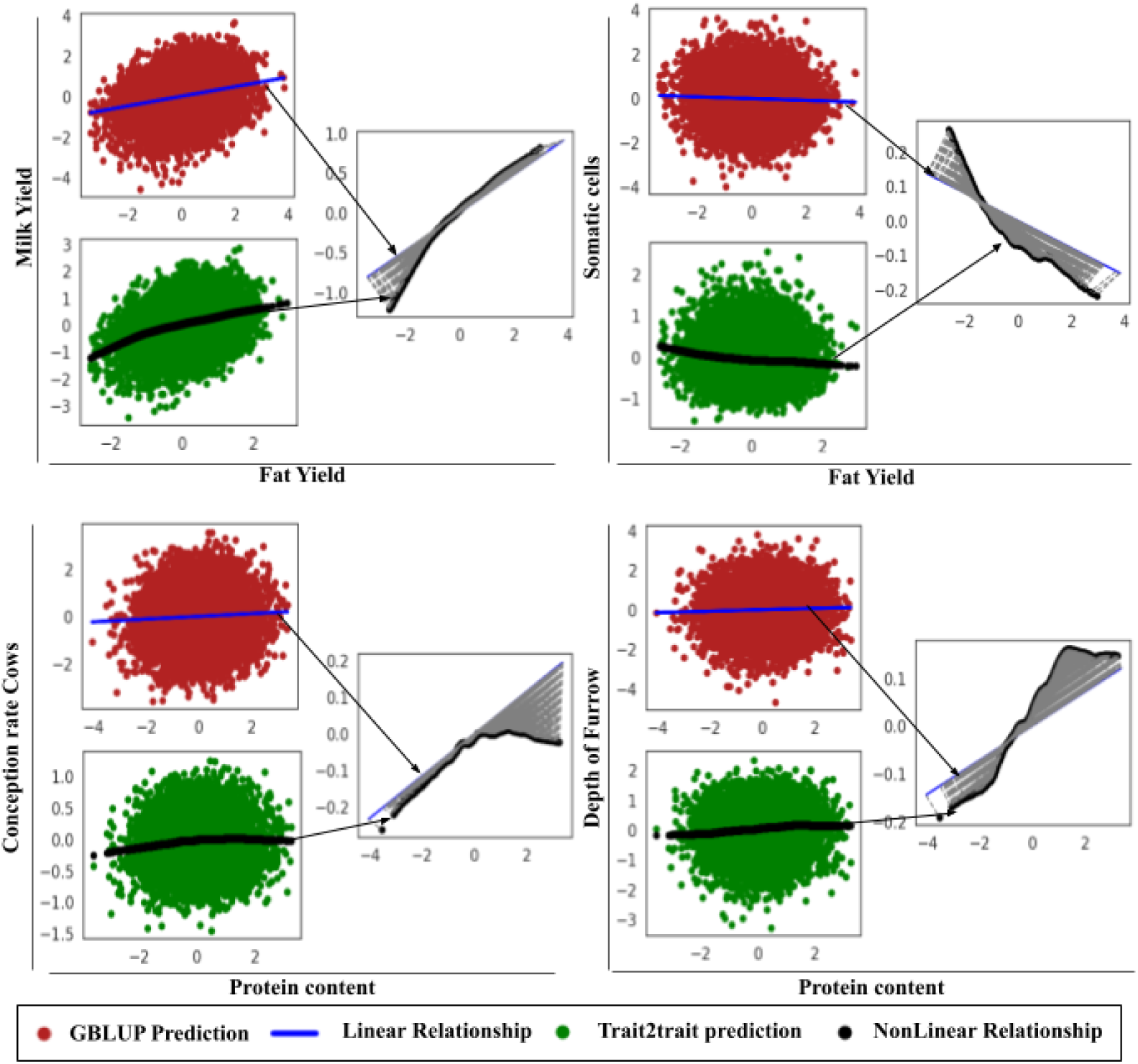
Plots of the predicted genetic values with GBLUP, DLGBLUP and the relationship representation between traits on real data. The red dots represent the predicted genetic value using GBLUP, in green the predicted genetic value using DLGBLUP, in black the predicted relationship representation using MeanTrait2Trait, and the blue line the linear relationship between two traits predicted using GBLUP. The traits in (y-axis) represents the dependent trait predicted from the trait in x axis (input) for MeanTrait2Trait model.

**Table 7:**
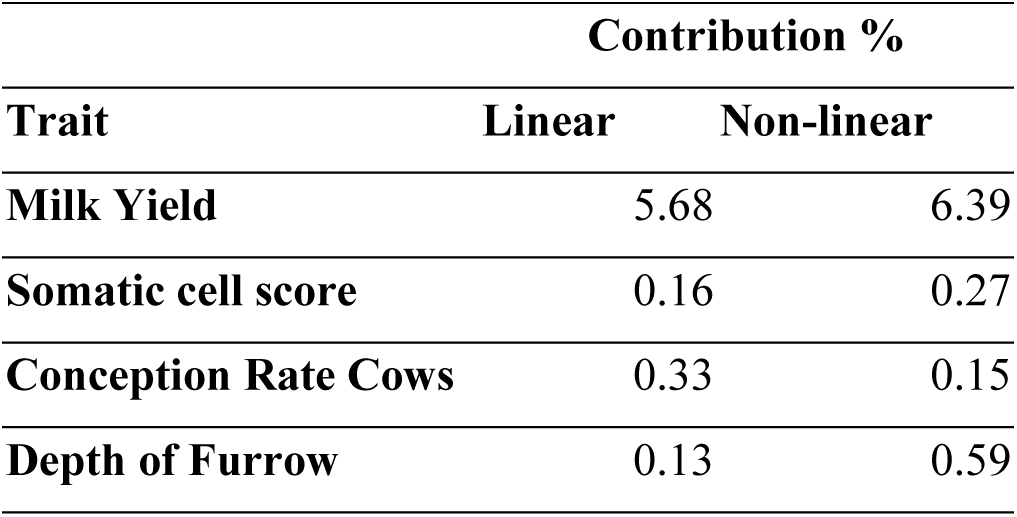
Contribution of linear and nonlinear relationship prediction for real data.

| Trait | Contribution % |  |
| --- | --- | --- |
|  | Linear | Non-linear |
| Milk Yield | 5.68 | 6.39 |
| Somatic cell score | 0.16 | 0.27 |
| Conception Rate Cows | 0.33 | 0.15 |
| Depth of Furrow | 0.13 | 0.59 |

## Discussion

In this study, we proposed a novel hybrid model, DLGBLUP, which integrates DL and statistical methods to perform a multi-trait prediction of nonlinearly related genetic values. Such nonlinear genetic relationships between traits have not been greatly explored in GP, neither using DL, nor statistical methods. In fact, DL methods in GP have been widely explored simply as an alternative for the statistical methods, and the results obtained with DL in real data sets have not yet significantly overcome those from GBLUP [9].

One possible reason why DL did not achieve higher performance in GP compared to GBLUP, may be the fact that in GP, the objective falls in predicting the additive GV, which are not directly observed in the phenotype data, without prior assumptions for DL models. While most studies implemented models that have been previously applied in different domains, in GP these models have to operate on the genotype-phenotype relationships - built-up in the hidden layers of DL with lack of biological interpretability - to predict the GV. Some studies incorporated a biological interpretation into their models, and others started to benefit from both DL and statistical methods, which performed better than those DL models used before [36, 25]. In this study, we proposed DLGBLUP, a model that takes GBLUP’s predictions and then adjusting the predictions according to the relationships between traits to predict a new GV.

Genomic selection in livestock or plant breeding programs consists of integrating the information of various traits into a SI, to select individuals that ensure genetic progress for traits of commercial interest, while not adversely impacting other traits that may be correlated to the targeted ones. To achieve this progress, two conditions are required: 1) A precise prediction of the individuals’ GV and 2) A good understanding of the relationships between traits. If one of these conditions is unavailable, accurately selecting animals for multiple traits may be challenging. Until now, statistical methods – and particularly GBLUP – have been the leading choice of breeders for performing genetic evaluations. On the prediction level, these statistical methods showed a competitive performance regarding their accuracy of prediction and current computational resources required. These methods are, however, restricted to the assumption of linearity for the relationships between traits. While this assumption may not affect the ranking of individuals on an intra-trait perspective, in the case of nonlinearity in the relationships between traits, the assumption of linearity may have an effect on the inter-trait level, when performing a genetic evaluation.

By accounting for the possibility of nonlinear relationships between traits, we have shown with our study that the DL and DLGBLUP models outperformed GBLUP on the simulated nonlinear traits, successfully capturing the shape of the relationship, especially if the provided input is more accurate. In addition, we demonstrated the critical role of an accurate SNPs selection step in minimizing input noise without eliminating important information, to enhance prediction accuracy. A significant advantage of DLGBLUP is its complementarity with other single or multi-trait statistical methods, such as GBLUP or even advanced approaches like the single-step method, which can predict values for non-genotyped individuals using pedigree information. This complementarity enables DLGBLUP to achieve superior results compared to using DL alone. Another motivation for this approach is its efficiency, requiring fewer parameters and less training time than a full DL model. This efficiency is achieved by eliminating the need for genotype data in the DL model, relying instead on predicted genetic values, which makes it feasible to implement using R, for example. The reason DLGBLUP outperformed the DL model might be that GBLUP is known for accurately predicting additive genetic values, ensuring the initial predictions (PGV_1) capture the additive effect with minimal errors. Unlike the DL SNP2Trait model that may face some issues like overfitting or insufficiently capturing additive genetic variance. So, starting with a precise and comprehensive PGV_1 from genomic data ensures accurate modeling and adjustment according to the full relationship between traits using the Trait2Trait module.

While research on the applications of DL for genomic prediction has become more and more widespread in the recent years, works that apply DL for multi-trait prediction models were less popular. Nevertheless, a previous study considered that relationships between the elements of the output layer could be learned and captured automatically by a neural network with shared neurons and weights [16]. However, this study did not explore whether these trait relationships are linear or nonlinear. In contrast, our study reveals that the SNP2Trait module for the DL model was unable to detect nonlinear relationships. Instead, the DL model required a proper and exclusive network to map one trait (input) to another (output), in order to capture nonlinearity. CNN was tested by including a convolutional block before the MLP block in the SNPs2Trait module. However, this archirecture underperform the MLP model (Table S3).

For the study on simulated data, the dependent traits were partially conditioned on the GV of a reference trait directly simulated from the genomic data. The relationships used in our study are over characterized. However, these examples are pertinent for a conceptual study aimed at developing a methodology. Demonstrating the model’s application in exaggerated nonlinear relationships enabled us to thoroughly test the method’s robustness.

In cattle, it may be possible to find non-linear genetic correlations between various traits, although these correlations have not been widely explored, given that usually BLUP is used as the evaluation model, limiting all between-trait correlations to be linear. For example, growth rate and feed efficiency might positively correlate up to a point, after which efficiency plateaus or declines. Fertility and milk production could exhibit a non-linear relationship, with moderate production levels having minimal impact on fertility, but very high levels reducing it. Longevity and production traits may positively correlate at moderate levels, while extreme production reduces lifespan. Mastitis and milking speed may also exhibit non-linear correlations, with both high and low speeds increasing mastitis risk due to tissue damage or prolonged exposure, respectively. These possibilities suggest that advanced methods are needed to fully understand and model these complex relationships for optimal breeding and management strategies.

On real data, our method was capable of detecting nonlinear relationships between multiple traits (Figure 8). This finding is important for selection and genetic progress, as we showed through simulation data. A good understanding of these relationships can be useful to manipulate the form as desired in the next generation and maintain a good genetic gain. However, this detection did not improve the prediction accuracy of these traits, which can be justified by the fact that these nonlinear relationships were not evident at the population level. Additionally, the contributions of both linear and nonlinear relationships were too small compared to those observed in the simulated data. Note that in our evaluation, we used yield deviations obtained from a previous pre-processing as the target, which can alter the actual form of the relationship between the traits and lead to these results. Moreover, the application of such models to raw performance data would need to consider additional factors, such as phenotypic records that may deviate from normality and multiple records for many production traits. The complexity of these models would exceed the scope of our study’s objectives, which were to test whether DL could identify non-linear correlations between traits and use this information to improve prediction accuracy. Further work is required to extend our approach to raw real data.

Better understanding how multiple traits involved in a breeding program are related is sure to improve the genetic progress obtained by artificial selection. Here, we maintained the use of a linear SI to select individuals, however considered the additive GV obtained with the different models (GBLUP, DL, and DLGBLUP) for comparison, and showed that for all traits after seven generations, genetic gains from SI based on the additive GV from either DL or DLGBLUP were equal or superior to the gains from SI based on the additive GV from GBLUP. We also found that the form of correlation between traits could be detected in the subsequent generation, which confirms the transmissibility of these traits. However, as selection continued over several generations, the correlations, especially for quadratic traits, tended to become more linear. This shift occurs because selection tends to favor a particular portion of the population—in this case, individuals exhibiting linear positive correlations between traits. This process results in a more homogeneous population with simpler, more linear trait relationships over time.

## Conclusions

In this study, we proposed a hybrid DLGBLUP model, that accounts for nonlinear genetic relationships between traits to predict their GV. From input SNP data and initial GBLUP predictions, that excel on predicting additive effects, the DL module in DLGBLUP model can adjust the prediction for potential nonlinear relationships between traits, when pertinent. Applied to simulated data, DLGBLUP was successful in improving the accuracy of the predicted GV in scenarios where nonlinear relationships between traits was present, in comparison to GBLUP. This greater prediction accuracy of the non-linearly related traits was due to the ability of DLGBLUP in correctly identifying the mean patterns of such relationships. Moreover, we showed that selection using a SI built based on the additive GV from either DL or DLGBLUP achieved greater genetic gain than selection using a SI built based on the additive GV from the traditional GBLUP.

## Author contributions

FS designed the deep learning models and the data simulation method, performed all the analysis, and did the main writing of the manuscript. PC participated in the conceptualization of the study, in the discussion and interpretation of the results, and in revising the manuscript. HG participated in the design of the deep learning models, and in revising the manuscript. RS participated in the discussion and interpretation of the results. TT contributed to the interpretation of preliminary results, leading to relevant decisions in the conception of the study. TMH participated in designing the data simulation method, in the design of the deep learning models, and in revising the manuscript. BCDC participated in the conceptualization of the study, in designing the data simulation method, in the discussion and interpretation of the results, and contributed to the writing of the manuscript. All authors read and approved the final manuscript.

## Acknowledgements

This work was supported by INRAE Metaprogramme DIGIT-BIO (Digital biology to explore and predict living organisms in their environment). We would like to think GenIALearn team for their valuable discussion and contribution, and Jocelyn de Goer De Herve from (INRAE /UMR EPIA) for managing the server on which DL work was done.

## Availability of data and materials

All source code in this study an dsimulated data are freely available at https://github.com/fshokor/DLGBLUP. The real data are not publicly available due to privacy or ethical restrictions.

