## Supplemental Figures for "Deep Learning and GBLUP Integration: An Approach that Identifies Nonlinear Genetic Relationships Between Traits"

**Additional files**

**Figure S1: Prediction and selection results for sinusoidal trait.**

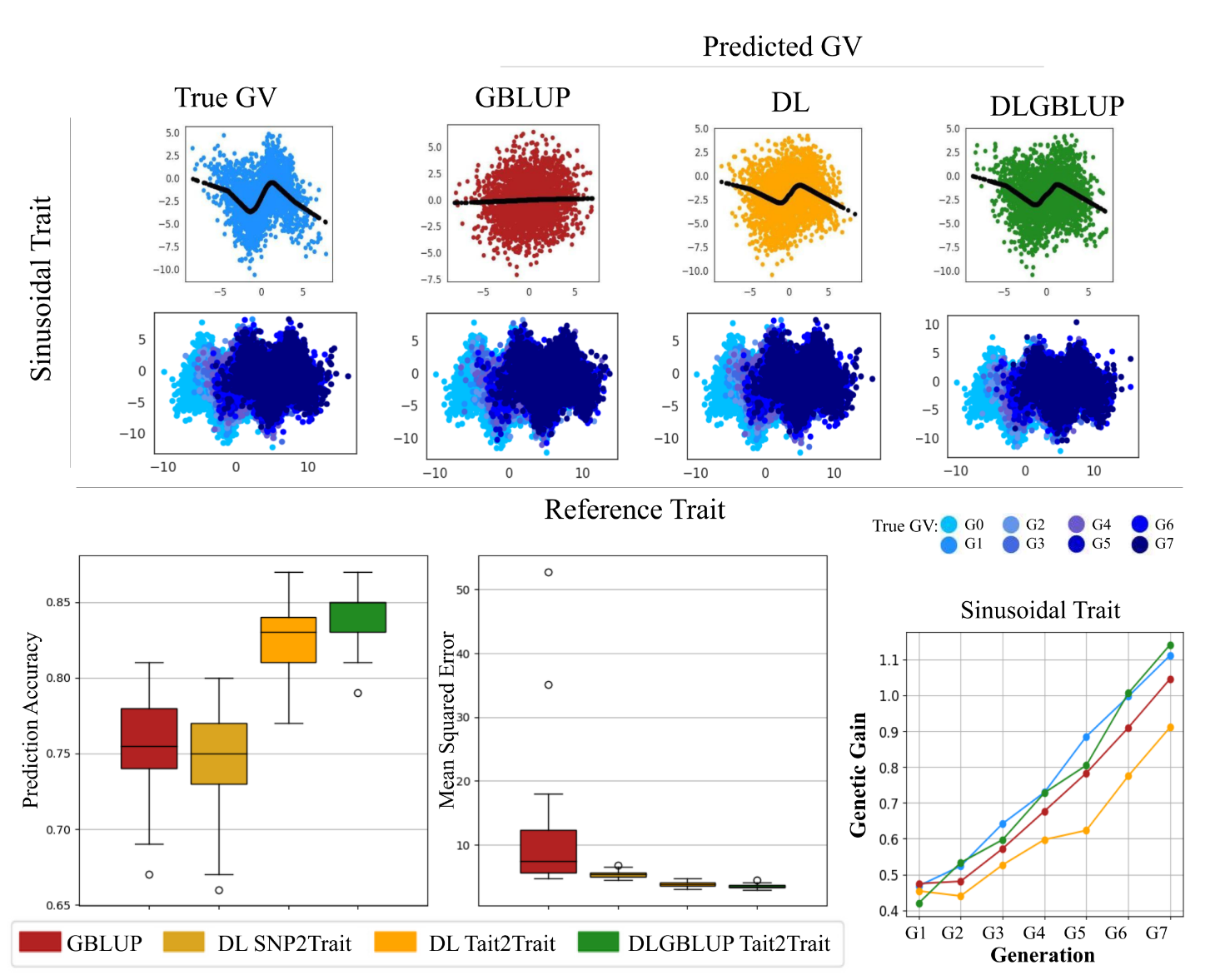

**Figure S2**: **Comparison plots of predicted genetic relationship between traits with different prediction model.**

Description: The reference trait is represented on the x-axis and all dependent traits on the y-axes. The colored points represent the PGV of GBLUP, DL SNPs2Trait, DL Trait2Trait and DLGBLUP and the TGV of a population. In black the mean relationship between traits as obtained from MeanTrait2Trait module. The prediction was made using genomic data with just QTL, h^2^ equal 0.3 and 50 specific QTLs.

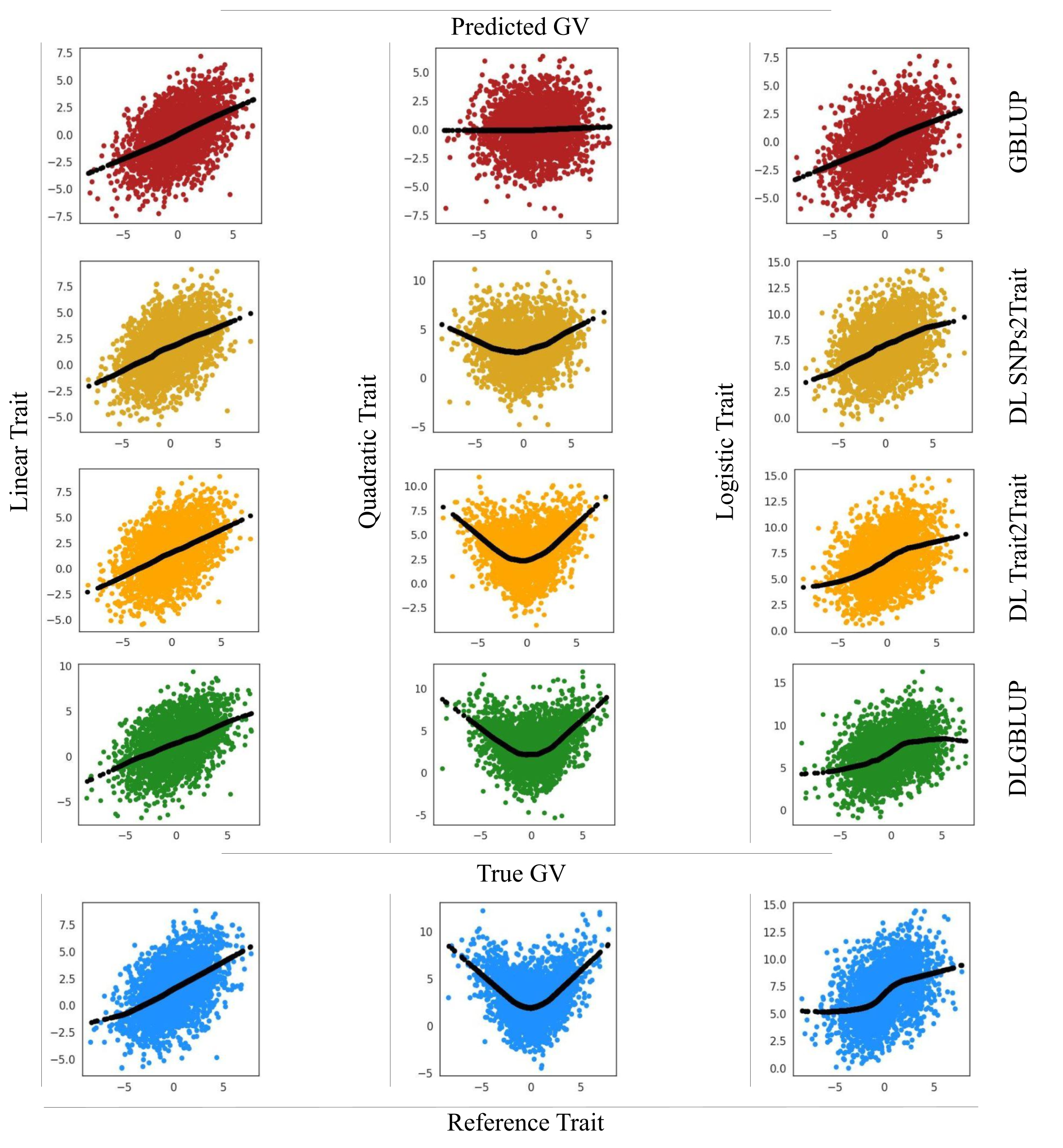

**Figure S3**: **Comparative plots of traits progression across 8 generations under selection.**

Description: Plots of relationship between the TGV of reference trait (x-axis) and dependent traits (y-axis) over 8 generations based on additive PGV of GBLUP, DL and DLGBLUP.

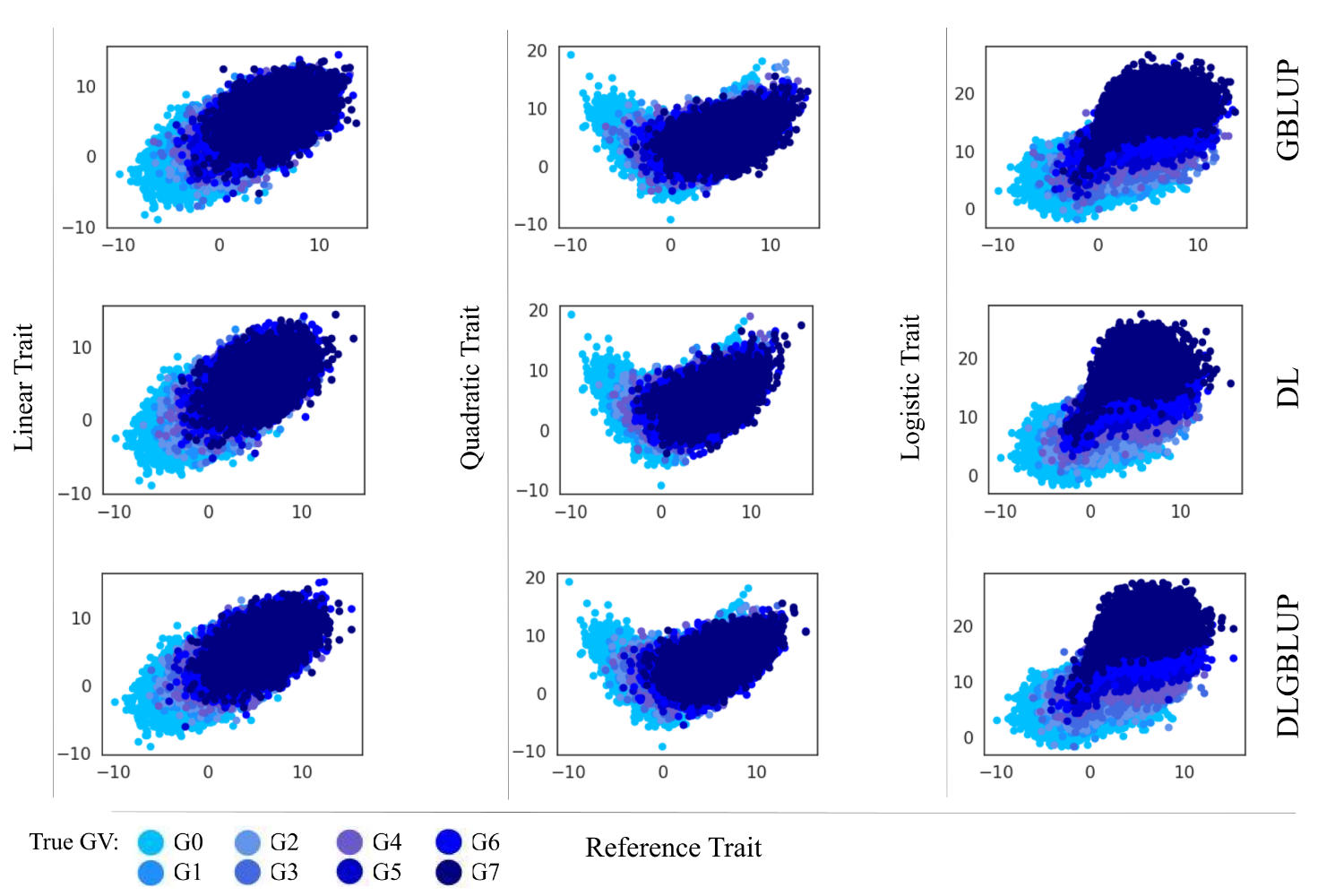
